# Sex differences in gene expression and proliferation are dependent on the epigenetic modifier HP1γ

**DOI:** 10.1101/563940

**Authors:** Pui-Pik Law, Ping-Kei Chan, Kirsten McEwen, Huihan Zhi, Bing Liang, Chie Naruse, Masahide Asano, Kian-Cheng Tan-Un, Godfrey Chi-Fung Chan, Richard Festenstein

## Abstract

Sex differences in growth rate in very early embryos have been recognized in a variety of mammals and attributed to sex-chromosome complement effects as they occur before overt sexual differentiation. We previously found that sex-chromosome complement, rather than sex hormones regulates heterochromatin-mediated silencing of a transgene and autosomal gene expression in mice. Here, sex dimorphism in proliferation was investigated. We confirm that male embryonic fibroblasts proliferate faster than female fibroblasts and show that this proliferation advantage is completely dependent upon heterochromatin protein 1 gamma (HP1γ). To determine whether this sex-regulatory effect of HP1γ was a more general phenomenon, we performed RNA sequencing on MEFs derived from males and females, with or without HP1γ. Strikingly, HP1γ was found to be crucial for regulating nearly all sexually dimorphic autosomal gene expression because deletion of the HP1γ gene in males abolished sex differences in autosomal gene expression. The identification of a key epigenetic modifier as central in defining gene expression differences between males and females has important implications for understanding physiological sex differences and sex bias in disease.

## Introduction

Sex dimorphism in physiology as well as gene expression pattern has been observed in many animal species including mice^1–6^ and humans^7–10^ during early development. For example, sex dimorphism in growth rate of mammalian embryos before gonadal sex differentiation has been shown by a number of previous studies, where males were found to be developmentally more advanced relative to females^2–4^. These differences are attributed to the difference in sex chromosome complement which are the only factors that differ between male (XY) and female (XX) zygotes and are thought to set up life-long sex differences. Indeed, using the ‘Four Core Genotypes’ mouse model, a number of studies have shown the crucial role of sex chromosome complement, independent of hormones in exerting divergent effects over metabolic and behavioural phenotypes in adults^11–13^. Further understanding of how sex dimorphism is regulated during early development is therefore important as sex chromosomes may differentially prime the genome and impact on developmental programming and other sexual differences (including disease susceptibility) later in life^14, 15^. However, little is known about the molecular mechanisms underlying these sexual differences.

Heterochromatin protein 1 (HP1) is a non-histone chromosomal protein that was initially discovered in *Drosophila* to be enriched at heterochromatic regions^16^. HP1 is highly conserved through evolution and its homologues are found in almost all eukaryotes except *Saccharomyces cerevisiae*^17^. In mammals, the three HP1 isoforms alpha (HP1α), beta (HP1β) and gamma (HP1γ) display a high degree of structural and biochemical similarity^17^ but appear to have distinct nuclear localisation^18–21^, expression profiles^22^, non-redundant functions^23–27^ and isoform-specific binding partners^28^. Among the three HP1 isoforms in mammals, HP1γ is distinguished by its unique subnuclear localisation pattern in both euchromatic and heterochromatic regions in interphase nuclei ^18, 21, 29, 30^.

HP1 has been regarded as a silencing protein with its interaction with the histone mark H3K9 methylation that associated predominately with silent heterochromatin^31–33^ as well as its ability to multimerise to interact with the H3K9 methyltransferase^34^. However, evidence accumulated in the last decade has implicated HP1 in a wide range of nuclear functions^35^. Of particular interest, is the implication that HP1γ participates in gene activation: (1) It has been shown that depletion of HP1γ in mammalian cells can result in down-regulation of gene expression^36, 37^, (2) HP1γ has been shown to associate with transcriptionally active genes^36, 38, 39^ and (3) interact with the RNA polymerase II ^38, 39^ and mediator components^36^. Yet, the exact nature and mechanisms whereby HP1γ contributes positively to gene expression is unclear.

Previously, it has been shown that depletion of a *Drosophila* HP1 homologue, HP1a, resulted in male-biased lethality^40^. In mice, HP1γ deletion resulted in growth retardation^41^, however, in that study both placental insufficiency and impaired proliferation were implicated as potential causes. Although defects in germ cell development have been identified in both sexes^42, 43^ in HP1γ deficient mice, another study showed that one female with a hypomorphic mutation on both alleles of HP1γ had normal ovaries^26^. These findings, together with our previous finding implicating HP1 in the regulation of sex-dimorphic autosomal gene expression in murine thymic cells^44^ raised the question as to whether HP1 was also contributing to sex dimorphism in early development of mice.

We directly addressed here whether HP1γ is involved in regulation of sex difference in early development. Using an HP1γ knockout mouse model, we found that the proliferation advantage of the primary embryonic fibroblasts derived from male E13.5 embryos is dependent upon the epigenetic modifier HP1γ. Strikingly, analysis of MEF’s transcriptome revealed that HP1γ is crucial for maintaining nearly all sex differences in autosomal gene expression by regulating the expression level of these genes in males.

## Results

### Sex dimorphism in cellular proliferation rate of primary mouse embryonic fibroblasts is dependent on HP1γ

In line with previous studies^2–4^, we observed a higher proliferation rate in male mouse embryonic fibroblasts (MEFs) derived from embryonic day 13.5 (E13.5) embryos than in female MEFs (Figure 1A). To investigate this sex difference further, we analysed the genome-wide transcriptional profile of these cells by RNA-sequencing (RNA-seq). Using Gene Set Enrichment Analysis (GSEA), we found a significant difference in the expression of a set of cell cycle related genes between males and females (Figure 1C and Figure S1A), consistent with the observed difference in cell proliferation rate.

**Figure 1.**
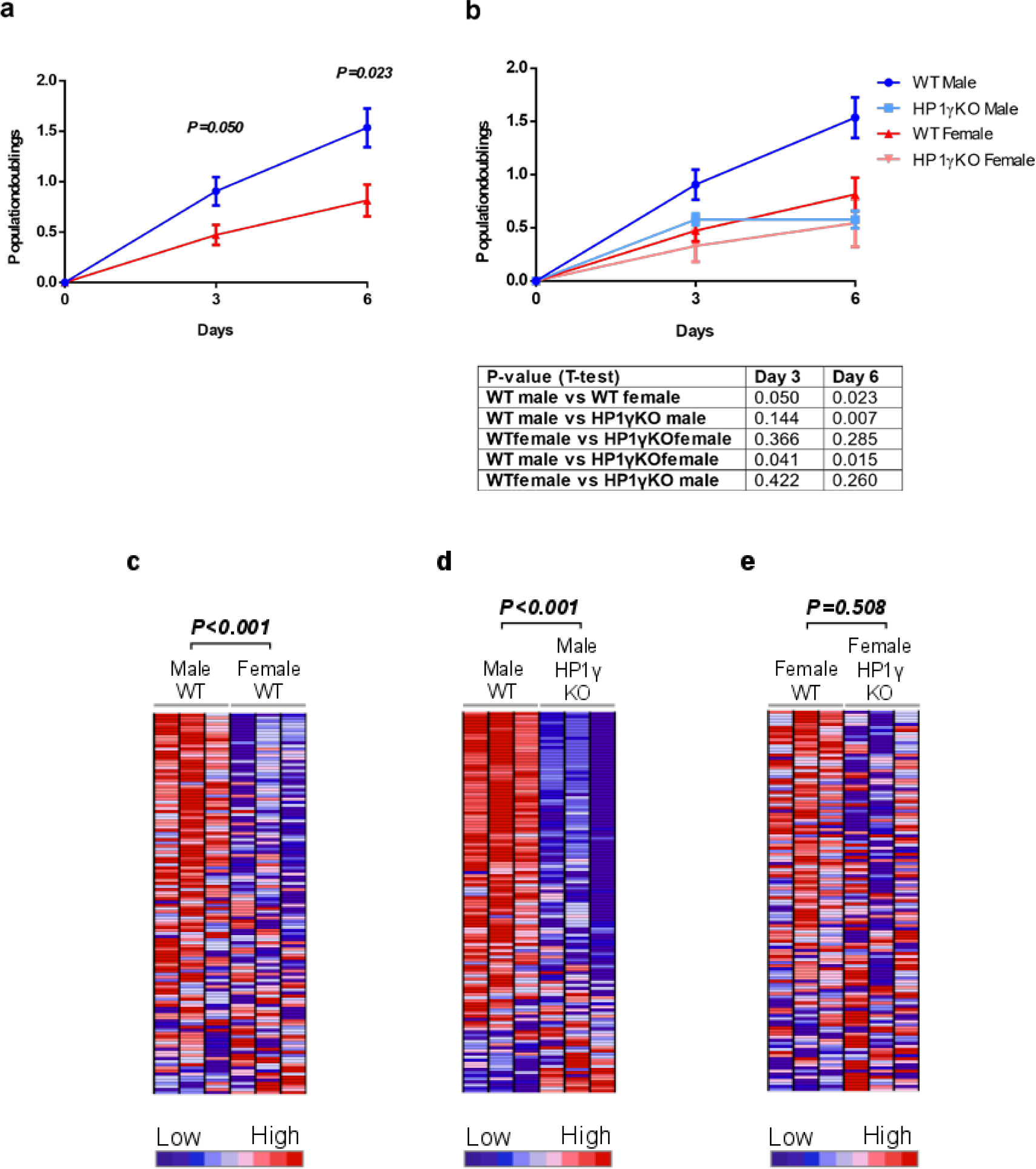
Sexually dimorphic proliferation rate of male and female E13.5 mouse embryonic fibroblasts (MEFs) is dependent on HP1γ. **a, b**, Growth curve of male and female MEFs with or without HP1γ showing population doublings from day 0 to day 6. n=8 for WT male; n=4 for HP1γ KO male; n=5 for WT female; n=3 for HP1γ KO female. Error bars: SEM. P-values for each time point were obtained by two-sided Student’s t-test. **c-e**, Genes with functions in cell cycle are affected by sex and HP1γ deletion. Gene set enrichment analysis (GSEA) of the transcriptome of male and female MEFs with or without HP1γ on cell cycle gene set. Comparisons were made between WT males and WT females (**c**), WT and HP1γ KO males (**d**) and WT and HP1γ KO females (**e**). The full list of genes shown on Fig. 1c to e can be found in extended Fig. 1a to c. Results were obtained from three MEF lines of each genotype derived from individual embryos. The data analysed here were derived from RNA-seq experiments that were repeated three times for each genotype.

Strikingly, upon removal of HP1γ (Figure 2), the normally higher proliferation rate of male wild type (WT) MEFs was reduced to a comparable level to that observed in WT females while the proliferation rate of female MEFs was largely unaffected (Figure 1B). GSEA of the transcriptomic data of these cells also showed a significant change in a set of cell cycle related genes in males but not in females upon HP1γ knockout (KO) (Figure 1D, 1E and Figure S1B, S1C). The difference in proliferation in WT male and female MEFs does not appear to be due to the difference in HP1γ level as a similar level of HP1γ gene expression was found in both WT male and female MEFs (Figure S3). Thus, depletion of HP1γ specifically affects proliferation rate of male MEFs and hence equalized the proliferation rate between the sexes. Thus, these data suggest that HP1γ is required for maintaining the sex difference in growth rate.

**Figure 2.**
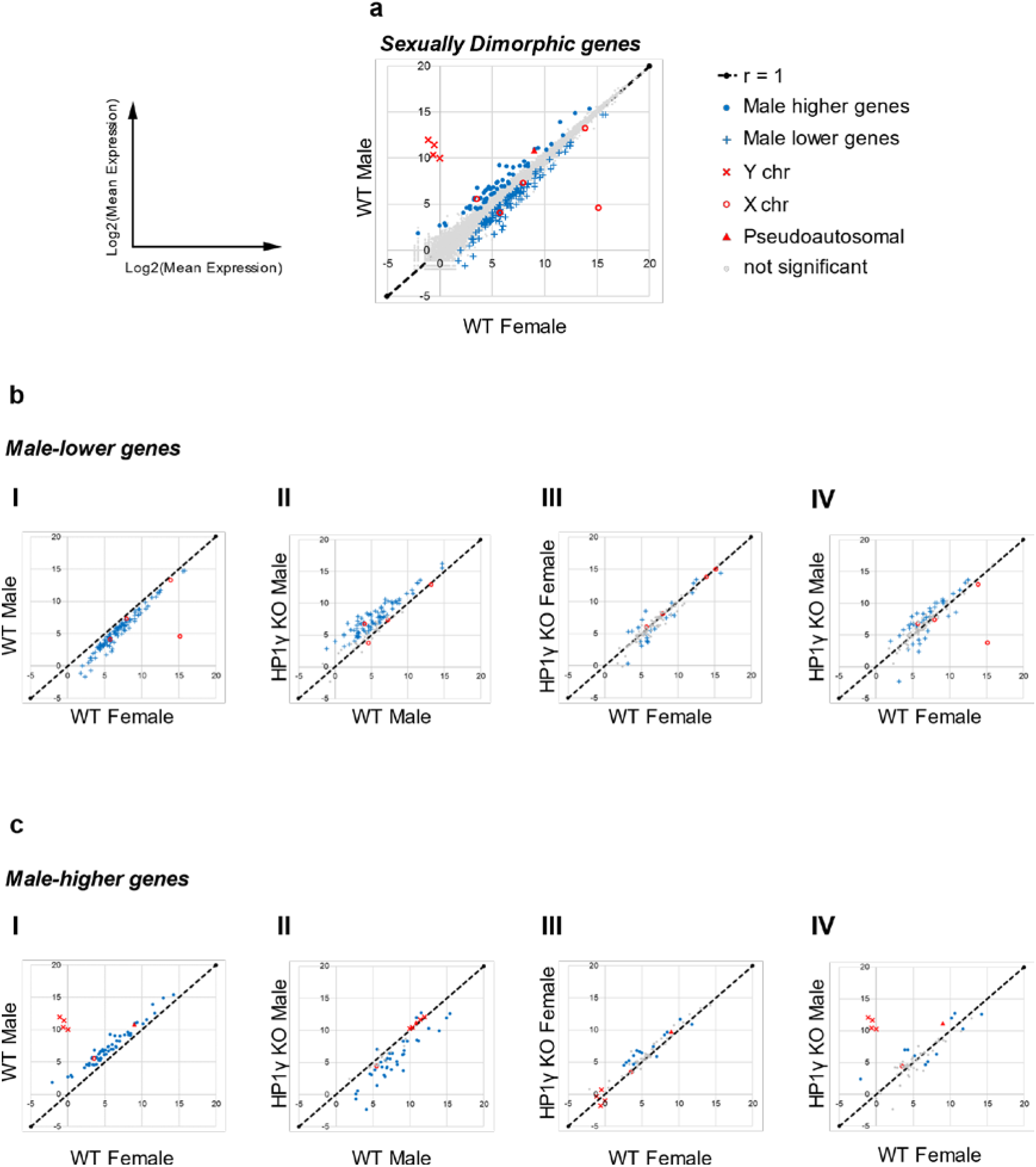
HP1γ regulates sexually dimorphic autosomal genes. **a**, Density plot of differentially expressed genes comparing WT male and female MEFs. 176 genes (out of 23420 genes) display sexual dimorphic gene expression. **b**, Density plot of 114 ‘male-lower genes’ which are repressed in WT males relative to WT females **(b(I))**. Upon HP1γ KO, most of the ‘male-lower genes’ were significantly upregulated in male (82 genes uperegulated; 0 gene downregulated; 32 genes not significantly changed) **(b(II)).** In female, most of these genes were not affected by HP1γ KO (7 genes uperegulated; 16 genes downregulated; 91 genes not significantly changed) **(b(III))**. Expression of ‘male-lower genes’ were also compared between HP1γ KO male and WT female MEFs (33 genes showed higher expression in males,10 genes showed higher expression in females and 67 genes not significantly different between two sexes exclusive of sex chromosome genes) (**(b(IV))**. **c**, Density plot of 62 ‘male-higher genes’ which are more highly expressed in WT males relative to WT females **(c(I))**. In males, most of the 62 ‘male-higher genes’ were downregulated in absence of HP1γ (3 genes uperegulated; 35 genes downregulated; 24 genes not significantly changed) **(c(II)).** A minimal effect was observed in females upon HP1γ KO (13 genes uperegulated; 2 genes downregulated; 47 genes not significantly changed) (**c(III)).** Expression of ‘male-higher genes’ were also compared between HP1γ KO male and WT female MEFs (6 genes showed higher expression in males; 5 genes showed higher expression in females; 45 genes not significantly different between two sexes exclusive of sex chromosome genes) **(c(IV)).** Results were obtained from three MEF lines of each genotype derived from individual embryos. Therefore, the data analysed here were derived from RNA-seq experiments that were repeated three times for each genotype. Genes showing ≥1.5-fold difference with P-value <0.05 between conditions compared were defined to be statistical significantly changed.

### HP1γ regulates sexual dimorphic expression of autosomal genes

The observed differences in response to HP1γ deficiency in males and females prompted us to ask how broad HP1γ’s role is in regulating sex-specific gene expression and hence contributing to the maintenance of sex differences. To address this question, sexually dimorphic genes (defined as genes showing ≥1.5-fold difference between WT males and females (P-value <0.05)) were identified from the transcriptomic data derived from the MEFs (Figure 2A and Table S2, S3) and tested for their sensitivity to HP1γ deficiency in both males and females. As well as genes encoded by the sex chromosomes, autosomal genes also showed dimorphic expression when WT males and females are compared (Figure 2A). These genes fall into two categories (1) 114 ‘male-lower genes’ that express at a lower level in WT males compared to females (Figure 2B(I)) and (2) 62 ‘male-higher genes’ that express at a higher level in WT males than females (Figure 2C(I))).

Strikingly, in males, the ‘male-lower genes’ were nearly all upregulated in the absence of HP1γ (Figure 2B(II)). In contrast, most of these genes were not changed in females upon HP1γ KO (Figure 2B(III)). These results suggest that, in the WT condition, HP1γ has an important role in repressing these genes in males and maintains sexually dimorphic gene expression. Indeed, direct comparison of the male HP1γ KO MEFs with the WT female MEFs revealed a marked loss of sex difference in the expression of these genes (Figure 2B(IV)). Almost all of the 62 ‘male-higher genes’ (Figure 2C(I)) were downregulated when HP1γ was knocked out (Figure 2C(II)), suggesting that in males most of these genes require HP1γ to keep them at a higher level than that seen in females. Comparison of the male HP1γ KO MEFs with the WT female MEFs revealed that HP1γ was required for the sex-difference in gene expression of these genes (Figure 2C(IV)). Similar to the ‘male-lower genes’, a minimal effect was observed on the ‘male-higher genes’ in females upon HP1γ KO (Figure 2C(III)). The changes in expression of a subset of the sexually dimorphic genes were confirmed by quantitative real-time PCR (qRT-PCR) (Figure S4). To test whether the observed dysregulation of these sexually dimorphic genes in response to HP1γ KO was an artifact of the analysis of a small group of genes (i.e. 176 genes out of 23420 genes), 176 randomly selected non-sexually dimorphic genes were also examined for their sensitivity to HP1γ deficiency (Figure S5A). In the absence of HP1γ, these genes were largely unaffected in both sexes (Figure S5B and S5C). These genes were also expressed at a similar level in HP1γ KO males compared to WT females (Figure S5D). Overall, these results implicate HP1γ in regulating the sexually dimorphic gene expression in MEFs and show that the effect is greater in male MEFs than female, suggesting an essential role for HP1γ in establishing or maintaining the difference in autosomal gene expression between the sexes.

### HP1γ deficiency leads to sex-biased dysregulation of a range of biological processes that are known to differ between the sexes

Our results revealed a male-specific defect in cell proliferation rate as well as dysregulation of sexually dimorphic gene expression in response to HP1γ deficiency. To investigate whether HP1γ may also affect other cellular processes in a sex-biased manner, genes that were specifically affected in males or females in response to HP1γ deficiency were identified (Figure 3A-3C) and the functional relevance of these genes was revealed using Gene Ontology (GO) analysis (Figure 4 and Tables S4 and S5). Consistent with previous studies, both up- and down-regulated genes were identified upon HP1γ KO in our system, implicating HP1γ in both the activation and repression of gene expression (Figure 3A and 3B)^36, 37^. A dramatic sexually-dimorphic transcriptional defect in response to HP1γ KO was observed (Figure 3A and 3B). 3827 genes were specifically dysregulated in males but only 720 genes in females by at least 1.5-fold (P<0.05) (Figure 3C). 466 genes were dysregulated in both males and females (Figure 3C). This sex-biased response to HP1γ deficiency is reminiscent of the previously reported difference between males and females upon depletion of HP1a in *Drosophila*^40^ suggesting evolutionary conservation. Gene Ontology (GO) analysis of genes regulated by HP1γ revealed a larger number of biological processes associated with genes affected in males (47) compared to females (20) (Figure 4 and Tables S4 and S5). These include several biological processes reported to differ between the sexes such as progression of cell cycle^2–4^ as well as immune system related functions^45^ (Figure 4 and Tables S4 and S5). Interestingly, ‘glucose homeostasis’ which is also known to differ between the sexes^46, 47^ is found in the female affected genes but not the male affected genes. Notably, as well as potentially regulating sex-bias in embryonic growth, cell-cycle related differences are known to be important for sex-dimorphism in cancer susceptibility^48^, moreover, response to infection and susceptibility to autoimmunity is well known to differ between the sexes.

**Figure 3.**
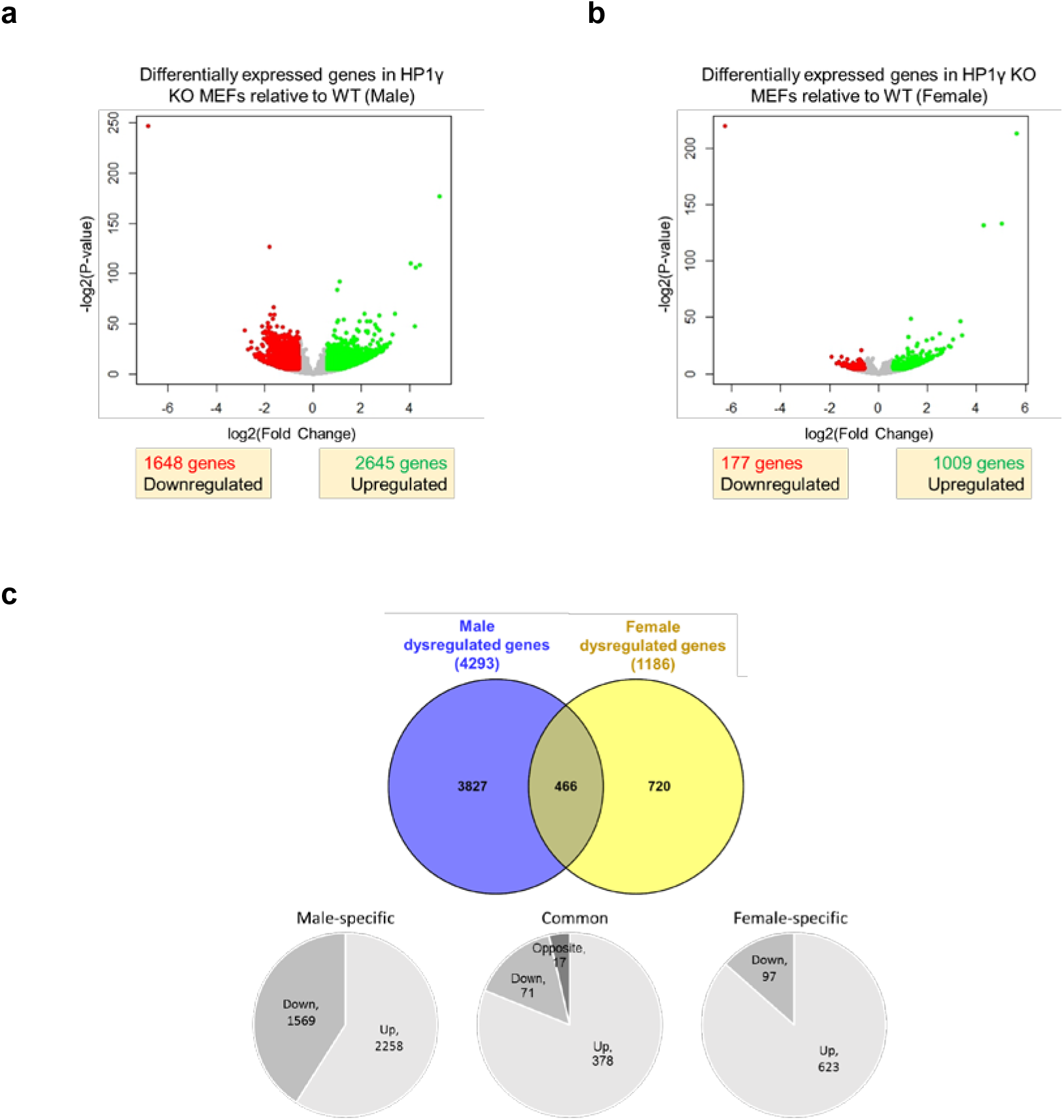
HP1γ-deficient MEFs display a male-biased transcriptional defect. Differentially expressed genes in male (**a**) and female (**b**) MEFs in response to knockout of HP1γ was revealed by RNA-seq. Results were obtained from three MEF lines of each genotype derived from individual embryos. (Fold change ≥1.5, P<0.05). **c**, Genes differentially expressed in male and female HP1γ-deficient MEFs and the number of genes up- or down-regulated or differentially expressed in opposite directions in each sex. The data analysed here were derived from RNA-seq experiments that were repeated three times for each genotype.

**Figure 4.**
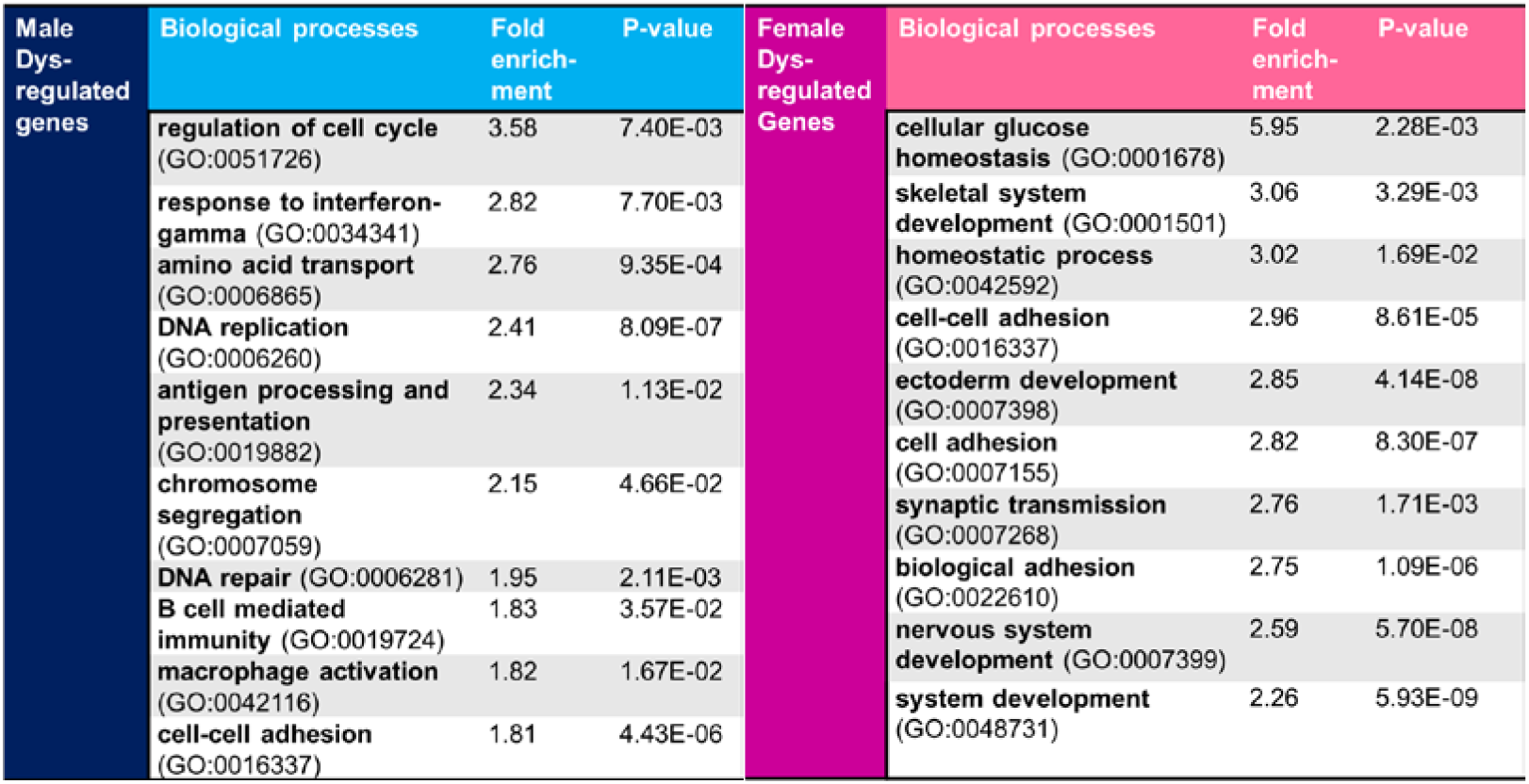
Sex-biased dysregulation of physiological functions upon HP1γ knockout. List of top 10 biological processes revealed to be associated with genes specifically dysregulated in males or females.

## Discussion

The results obtained in the current study suggest a male-specific role for HP1γ in regulating sexually dimorphic gene expression which in turn contributes to sexual differences. This is perhaps a surprising finding because HP1γ was found to be expressed at a comparable level in males and females (Figure S3). Interestingly, a study in *Drosophila* revealed male sex-bias in the effect on gene regulation upon depletion of the HP1 homolog HP1a ^40^ although that study did not examine the effect on sex dimorphic gene expression. Taken together with our results these findings might suggest evolutionary conservation of sex-specific effects of HP1 on gene regulation.

How the male-specific regulation of the sexually dimorphic genes is mediated by HP1γ remains to be further studied. In this study, sex-specific effects of HP1γ were studied in mouse embryonic fibroblasts derived from embryos at E13.5 whose fetal Leydig cells have just started to produce androgen^51^. It is therefore possible that the very low levels of testosterone found in these male embryos might contribute to the targeting of HP1γ in a sex-specific manner. However, regulation of pituitary growth hormone secretion by gonadal hormones before puberty in mice was shown to be minimal^52^. Thus, sexual difference observed here is likely to be regulated by sex chromosome complement rather than just sex hormones^53^. It is tempting to speculate that HP1γ may interact directly or indirectly with one or more factors contributed by the sex chromosome complement to give rise to the sex-specific effect observed.

The Y chromosome which is uniquely present in males may encode factors that contribute to HP1γ’s male-specific regulation of the ‘male-higher’ genes in males. It has been shown previously that HP1γ associates with the protein encoded by the *Sry* gene, which is crucial for male sexual development^54^ through indirect interaction with KAP1 and KRAB-O protein^55^. However, most of the Y chromosome genes, including *Sry*, are largely repressed in MEFs with only four of them showing pronounced expression in our transcriptomic data. Two genes were of particular interest, *Uty* and *Kdm5d* which encode H3K27me3 and H3K4me3 demethylases respectively^56, 57^ (Table S7).

Although dosage difference of X chromosomes between males and females is compensated by X chromosome inactivation in female cells^58^, a number of X-linked genes were found to escape X-inactivation^59^. Indeed, we observed in our transcriptomic data female-biased expression of several known X-escapees^59^ (*Kdm5c*, *Eif2s3x* and *Kdm6a*) as well as some other X-linked genes which have not been previously reported as escaping X-inactivation (*5530601H04Rik*, *2900056M20Rik* and *Ppef1*) (Table S7). To contribute to the male-specific regulation of the ‘male-higher’ genes, the proteins encoded by these X-escapees may compete with HP1γ for binding to these genes in females to protect them from the activating effect of HP1γ.

Interestingly, when looking at the list of sex chromosome genes that are differentially expressed in males and females, we found two X-linked genes (*Ppef1* and 2900056*M20Rik*) which show female-biased expression and were upregulated in response to HP1γ knockout in males. Importantly, upon removal of HP1γ in males, the difference in expression level of these genes between males (KO) and females (WT) became insignificant (Table S8). *Ppef1* encodes a serine/threonine-protein phosphatase^60^. Although this protein has not been shown to regulate gene expression directly, being a protein phosphatase, it is possible that it regulates gene expression indirectly by activating or deactivating other protein factors. *2900056M20Rik*, also known as *Kantr* (Kdm5c adjacent non-coding transcript) which encodes a long non-coding RNA. It has been shown previously that deletion of *2900056M20Rik* results in an increase in expression of sets of genes involved in neuronal development and various cell signaling pathways and in murine embryonic and adult brain^61^. It is possible that the higher expression level of these genes in females is responsible for keeping the higher expression of these ‘male-lower’ genes in females compared to males. Removal of HP1γ may upregulate expression of *Ppef1* and *2900056M20Rik* in males and lead to upregulation of the ‘male-lower’ genes in males and normalization of sex difference.

Finally, it has been hypothesised that the inactive X-chromosome in females that is decorated with H3K9me3^62^ might be acting as a heterochromatic ‘sink’^63^ thereby sequestering HP1γ away from the euchromatic regions. Such an effect would operate in females effectively reducing the availability of HP1

In summary, we have identified the epigenetic modifier HP1γ as an essential component in the regulation of autosomal gene expression differences between the sexes which is likely to be important for understanding sex dimorphisms in physiology and disease.

## Acknowledgements

This work was supported by the Medical Research Council (MRC) (grant number MR/J007943/1). P.P.L was supported by the Joint University of Hong Kong-Imperial College London Ph.D. program. We thank the Medical Research Council London Institute of Medical Sciences (MRC LMS) Genomics Facility for Illumina sequencing services. Thanks to Prof. Christian Speck and Prof. Petra Hajkova for critical review of the manuscript.

