## Supplementary material for "Sex differences in gene expression and proliferation are dependent on the epigenetic modifier HP1γ": Methods

**Supplemental experimental procedures**

**Experimental mice**

All mice used in this study were maintained and handled according to the Imperial College Subcommittee for Animal Research guidelines and the British Home Office regulations. HP1γ-knockout mice were generated by a gene trap method where a single gene-trap retroviral vector (ROSAN β-geo) was inserted into intron 1 (998bp downstream of exon 1) of the *Cbx3* gene (encoding HP1γ)^1^ and backcrossed onto CBA/Ca background >10 times.

**Generation and maintenance of mouse embryonic fibroblasts (MEFs)**

WT and HP1γ homozygous knockout mouse embryonic fibroblasts (MEFs) were generated by breeding male and female HP1γ heterozygous knockout mice. E13.5 embryos were obtained and the heads and internal organs were removed. The body of individual embryo was finely minced in 1ml phosphate buffered saline (PBS) and incubated with 1.5 ml of 0.25% trypsin-EDTA (Sigma) at 37°C for 1.5min. 20ml of culture medium (Dulbecco’s modified Eagle’s medium (DMEM) supplemented with fetal bovine serum (FBS) (15% v/v), penicillin/streptomycin (Gibco®) (1% v/v), GlutaMAXTM-I 100x (Gibco®) (1% v/v)) was then added to the sample and the sample was passed through a 0.7µm cell strainer. Cells in the suspension were then collected by centrifugation at 300 x g at room temperature for 5min. The cells were then cultured in DMEM medium with supplements as specified above.

**Proliferation assay**

On day 0, 1x10^5^ of MEF cells at passage 1 obtained from individual embryos were seeded in each well of a 12-well tissue culture plate and incubated at 37°C, 5% CO_2_ for 3 days. Cells obtained from each embryo were treated as an individual biological replicate. On day 3, MEF cells in each well were typsinised and the cell numbers were determined using Guava Viacount reagent (Millipore) and the Muse® Cell Analyzer (Millipore). 1x10^5^ cells from each MEF cell line were then obtained and seeded in a fresh well of a 12-well tissue culture plate and incubated at 37°C, 5% CO_2_ for another 3 days. On day 6, cell numbers were determined again as stated above. Growth curves of these MEF lines were presented as population doublings which represents the total number of times the cells in the population have doubled since passage 1 (Day 0). This is calculated as:

Population doubling in each passage (Px) = log2 (Nx/105)

Accumulative population doubling = Sum (P1: Px)

**RNA extraction and complementary DNA (cDNA) synthesis**

Total RNA from MEFs was extracted using TRIZOL® Reagent (Invitrogen) following the manufacturer’s protocol. RNA extracted was treated with DNaseI using Ambion® DNA free kit (Invitrogen) following manufacturer’s instructions. RNA extracted was reverse transcribed into complementary DNA (cDNA) using the ThermoScript® (Invitrogen) reverse transcriptase and random hexamers following manufacturer’s protocol. cDNA generated was then analysed by qRT-PCR.

**Quantitative real-time PCR (qRT-PCR)**

Quantitative real-time polymerase chain reaction (qRT-PCR) was used to analyse samples from ChIP experiments and mRNA expression level. qRT-PCR was carried out using an Chromo4^TM^ system (Bio-Rad) with SYBR® Green JumpStart™ Taq ReadyMix™ (Sigma). mRNA expression levels are expressed relative to the expression of 18S rRNA and were analysed using the 2^–ΔΔCT^ method.

**RNA sequencing library preparation and sequencing**

RNA extracted from each MEFs line derived from individual embryos from different pregnancies were used to prepare each RNA sequencing library. RNA sequencing libraries were prepared using the TruSeq® RNA sample preparation kit v2 (Illumina®) following ‘Low Sample (LS) protocol’ provided by the manufacturer. 0.5μg of good quality RNA checked by Agilent 2100 Bioanalyzer (Agilent Technologies, Inc.) using the RNA 6000 Nano kit (#5067-1511, Agilent Technologies) was used per RNA-seq library preparation reaction. Prepared libraries were validated by Agilent 2100 Bioanalyzer using High Sensitivity DNA Analysis Kits (#5067-4626, Agilent Technologies, Inc.) following the manufacturer’s protocol. Samples of sufficient quality were sequenced using the HiSeq2500 (Illumina®) platform.

**RNA sequencing analysis**

Raw data obtained from sequencing were processed using RTA (ver. 1.17.21.3) with default filter and quality settings from Illumina®. The reads were demultiplexed with CASAVA 1.8.2 (allowing 0 and 1 mismatch). Raw reads (fastq files) were aligned to the mouse genome (mm10) using Tophat aligner (tophat2 ver. 2.0.11)^2^. Aligned read counts data were generated by HTseq work package (ver. 0.5.4p3). Differential expression analysis was performed using DESeq2 Bioconductor package^3^. Raw p-values were adjusted for multiple testing using the Benjamini-Hochberg procedure. Gene ontology (GO) statistical overrepresentation test was performed using the Panther classification system (PANTHER version 10.0)(http://pantherdb.org/) ^4^. Gene sets for the Gene set enrichment analysis (GSEA) was significant if normalised enrichment score >1.5, P<0.05 by absolute weighted signal-to-noise enrichment ^5^. Cell cycle gene sets were downloaded from KEGG (cell cycle genes from KEGG pathway mmu04110). Venn diagrams presented were generated by Venny 2.0.2 (Oliveros, J.C. (2007-2015). (<http://bioinfogp.cnb.csic.es/tools/venny/index.htm)>)

**Statistics**

Three biological replicates per mouse genotype were analysed for RNA-seq with no prior knowledge of variability. No further mouse sample size calculations proved necessary. Mice were analysed as the correct genotypes became available without any other form of selection. No formal randomization protocol was done. For Q-RTPCR validation two-sided student’s t test was used on three biological replicates per genotype using GraphPad Prism 6 software.

For the proliferation assays the researcher counting cells was blinded to the genotype.

**Data**

The RNA-seq data have been deposited in Gene Expression Omnibus (GEO) and given the accession number: TBC
