## Supplemental Figures for "Sex differences in gene expression and proliferation are dependent on the epigenetic modifier HP1γ"

**Supplemental Figure 1**

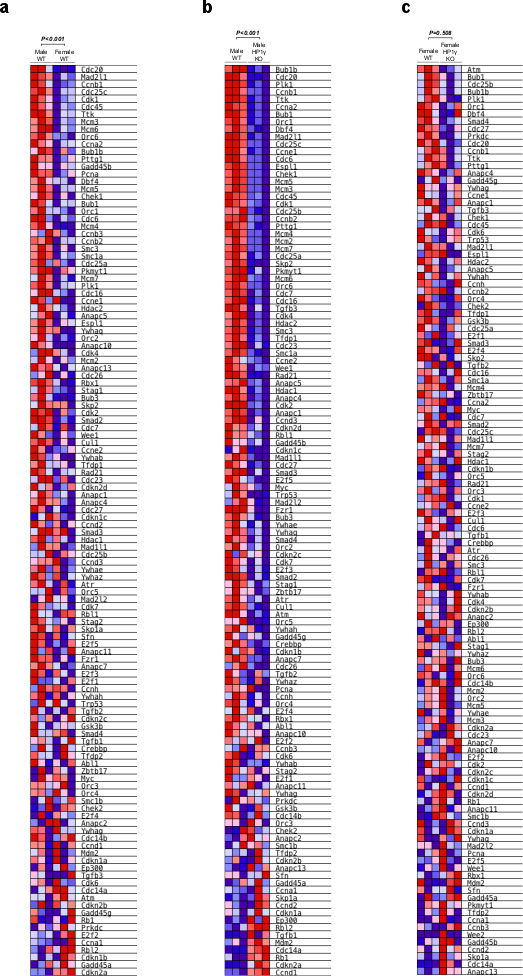

**Figure S1. a-c,** Extended heat map of genes with functions in cell cycle affected by sex and HP1γ deletion as presented in Figure 1C to 1E. Gene set enrichment analysis (GSEA) of the transcriptome of male and female MEFs with or without HP1γ on cell cycle gene set. Comparisons were made between WT males and WT females (**a**), WT and HP1γ KO males (**b**) and WT and HP1γ KO females (**c**). Results were obtained from three MEF lines of each genotype derived from individual embryos. The data analysed here were derived from RNA-seq experiments that were repeated three times for each genotype.

**Supplemental Figure 2**

**a**

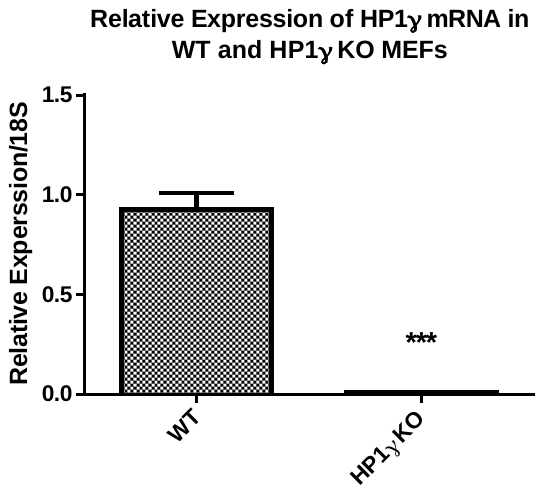

**b**

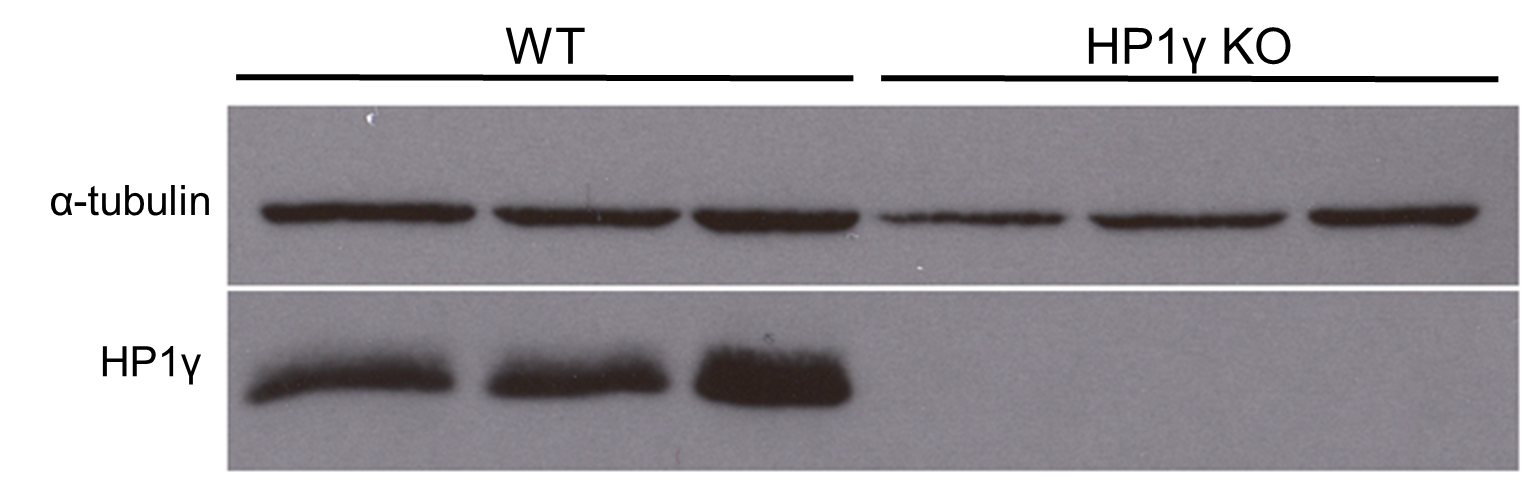

**Figure S2. Expression of HP1γ in WT and HP1γ KO MEFs.** **a,** Relative mean expression of HP1γ mRNA in MEFs derived from WT and HP1γ KO E13.5 embryos. Student’s t-test: ***P<0.0001 (compared to WT after being normalised against 18S expression). Error bars: SEM, n =16 for WT; 23 for HP1γ KO MEFs. **b,** HP1γ protein level in WT and HP1γ KO MEFs as shown by Western blot analysis. 3 representative experiments from MEFs derived from three 3 individual embryos are shown here.

**Supplemental Figure 3**

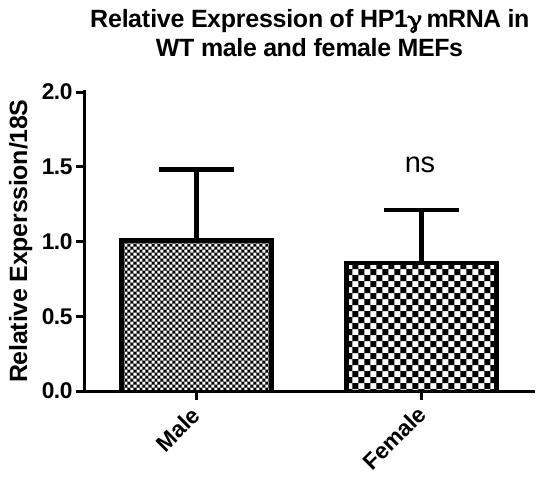

**Figure S3. Expression of HP1γ mRNA in WT male and female MEFs.** Relative mean expression of HP1γ mRNA in MEFs derived from WT male and female E13.5 embryos. Student’s t-test: ns = not significant (compared to male after being normalised against 18S expression). Error bars: SD, n=12 for male; n=10 for female.

**Supplemental Figure 4**

**a**	**b**

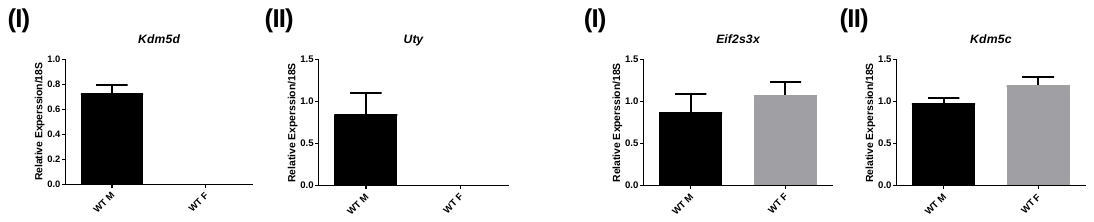

**c**

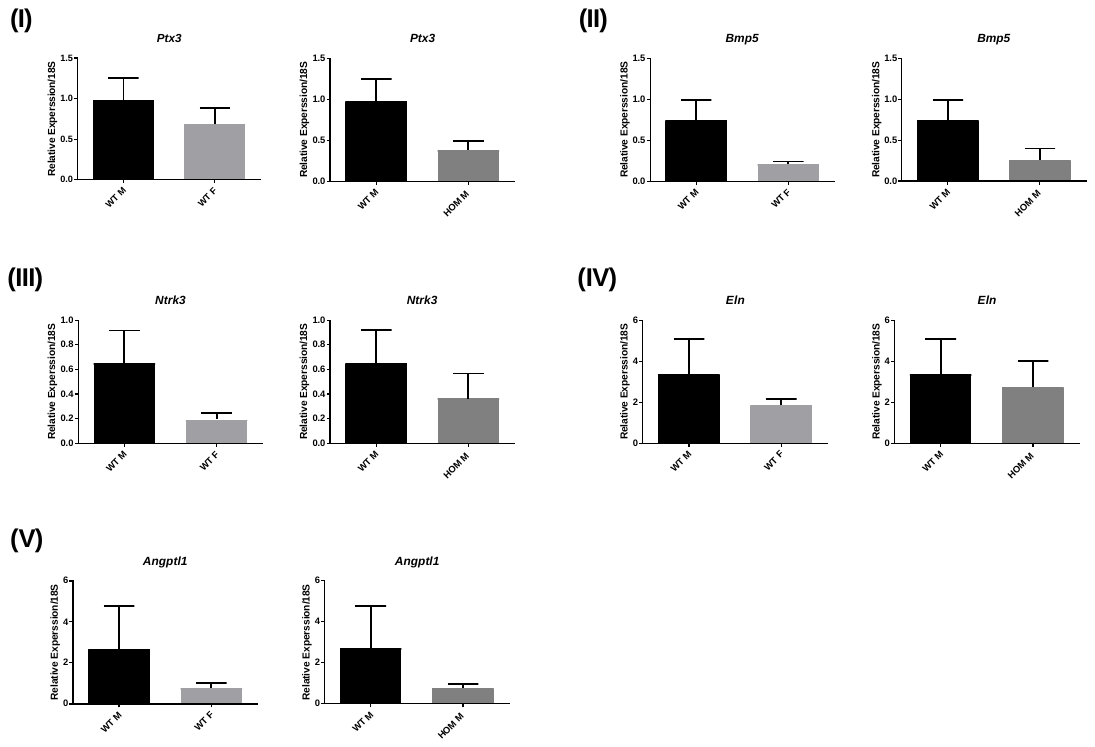

**Supplemental Figure 4 (cont.)**

**d**

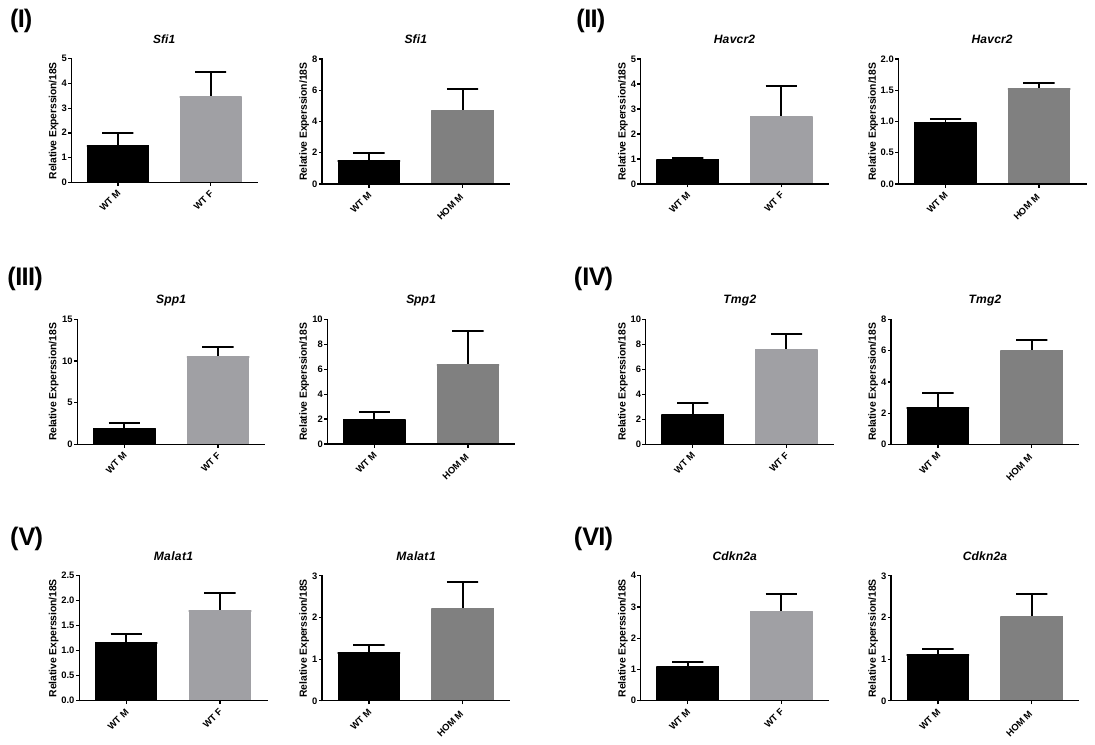

**Figure S4. Validation of RNA-seq result by qRT-PCR. a-d,** Relative expression of a subset of randomly selected Y chromosome genes (**a**), X chromosome genes (**b**), genes with relatively higher expression in males (**c**) and genes with relatively higher expression in female (**d**). Results were obtained from three MEF lines of each genotype derived from individual embryos. The data analysed here were derived from Q-RTPCR experiments that were repeated three times for each genotype. Error bar = SD.

**Supplemental Figure 5**

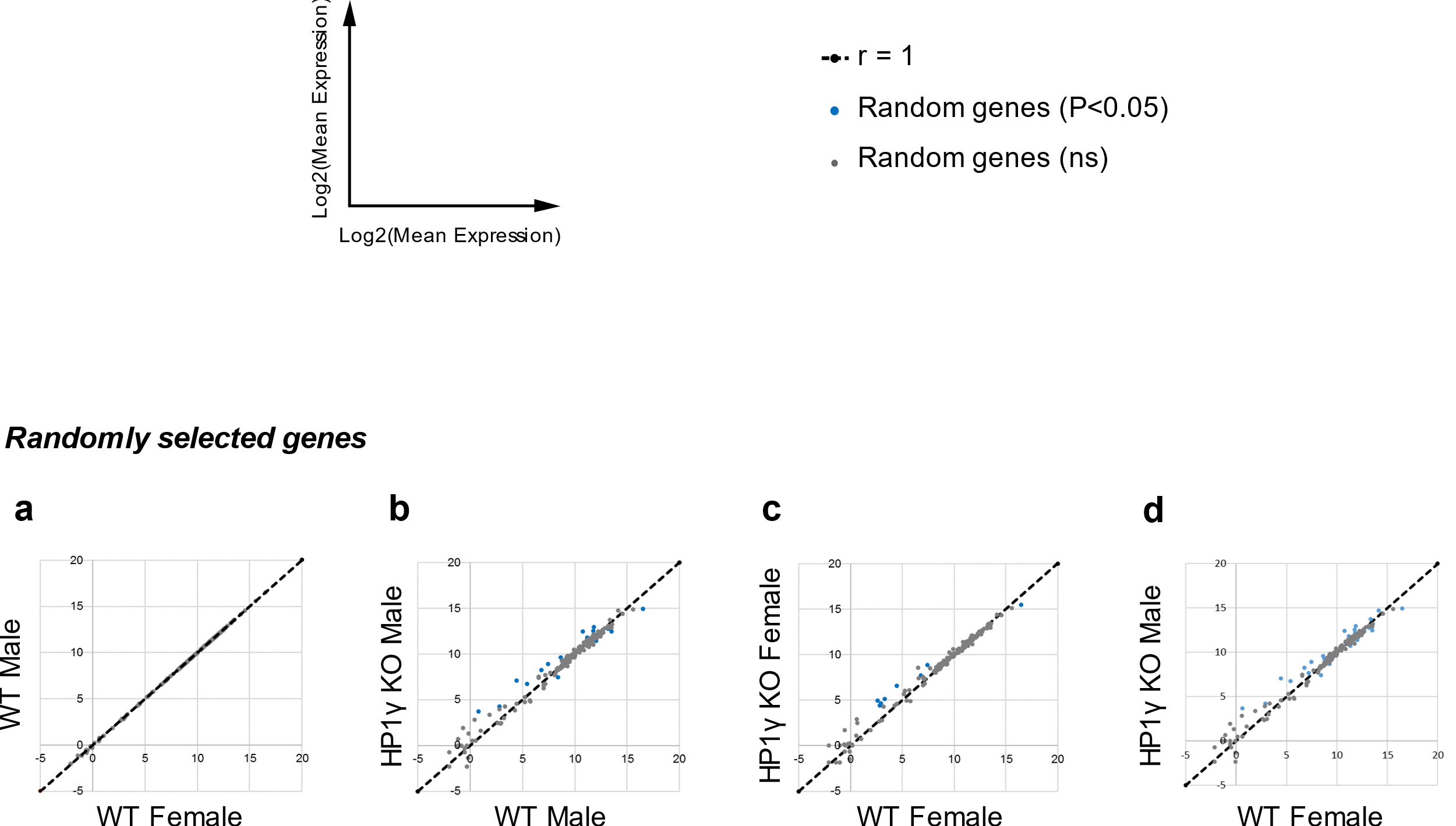

**Figure S5. Enrichment of HP1γ sensitivity was not observed on the 176 randomly selected non-sexually dimorphic genes in both males and females**. **a,** Density plot of 176 randomly selected non-sexually dimorphic genes. **b-d,** Expression of these genes were compared between WT and HP1γ KO male (**b**), WT and HP1γ KO female (**c**) and WT female and HP1γ KO male (**d**). No enrichment of HP1γ sensitivity could be observed on these genes in all comparisons.

**Table S1. List of primers used in this study**

| **Primer name** | **Gene target** | **Primer sequences (5’-3’)** |
| --- | --- | --- |
| Angptl1(F) | *Angptl1* | ATAGGTGGCCCAGCAAACTC |
| Angptl1(R) | *Angptl1* | CCACAGCGTCTTTGTTGTCTG |
| Bmp5(F) | *Bmp5* | AGCACCAGAAGGGTATGCTG |
| Bmp5(R) | *Bmp5* | ACATCAGGTGTACCAGGGTCT |
| HP1γ(F) | *Cbx3* | GGTCCAGGTCAGCCAGTCTA |
| HP1γ(R) | *Cbx3* | CCAGCCACGATTCTATTTCC |
| Cdkn2a(F) | *Cdkn2a* | GAAGGCTTCCTGGACACGC |
| Cdkn2a(R) | *Cdkn2a* | TGAGCTGAAGCTATGCCCG |
| Eif2s3x(F) | *Eif2s3x* | TTCTCTCCGAGCAAGATGGC |
| Eif2s3x(R) | *Eif2s3x* | GGTGTCAACTTGGTAACATCCAAT |
| Eln(F) | *Eln* | GAGTCTCGACAGGTGCTGTG |
| Eln(R) | *Eln* | GAGCCTTGGCTTTGACTCCT |
| Havcr2(F) | *Havcr2* | TGACCATGGGACCTACTGCT |
| Havcr2(R) | *Havcr2* | TCTCTCCGTGGTTAGGGTTCT |
| Kdm5c(F) | *Kdm5c* | GTACCAGTCTGGAGCCAACC |
| Kdm5c(R) | *Kdm5c* | GGTTCCGGATCAGGCTGTAG |
| Kdm5d(F) | *Kdm5d* | TTCGGTCCCACTACGAACG |
| Kdm5d(R) | *Kdm5d* | TTCCTCGTCTACTGTAGCAACT |
| Malat1(F) | *Malat1* | GGGGGAATGGGGGCAAAATA |
| Malat1(R) | *Malat1* | AACTACCAGCAATTCCGCCA |
| Ntrk3(F) | *Ntrk3* | CGGGAATTGAGACTGGAGCA |
| Ntrk3(R) | *Ntrk3* | GTTGGGAGCCATCAGCACT |
| Ptx3(F) | *Ptx3* | CTGCCCGCAGGTTGTGAAA |
| Ptx3(R) | *Ptx3* | ACCAACACTAGGGACTGGGA |
| Sfi1(F) | *Sfi1* | CGATGGCAAGAACAGCTTCTAA |
| Sfi1(R) | *Sfi1* | ACAGCCATCTCATTCTGCCAT |
| Spp1(F) | *Spp1* | CTCCTTGCGCCACAGAATG |
| Spp1(R) | *Spp1* | TTGTGTACTAGCAGTGACGGT |
| Tgm2(F) | *Tgm2* | ACAAGTACCCAGAGGGGTCA |
| Tgm2(R) | *Tgm2* | AGAGCAGGAGACGACACTCT |
| Uty(F) | *Uty* | TCCAAGACCACCAACTTCGC |
| Uty(R) | *Uty* | CCAAGATCTAACTTAAGAGCTCCAG |
| 18S (F) | Rn18s | ATGGTAGTCGCCGTGCCTAC |
| 18S (R) | Rn18s | CCGGAATCGAACCCTGATT |

**Table S2. List of male-higher genes showing ≥1.5-fold higher expression in WT males compared to WT females (P-value <0.05)**

|  | **Genes** | **Chromosome** |
| --- | --- | --- |
| 1. | *Eif2s3y* | Y |
| 2. | *Ddx3y* | Y |
| 3. | *Kdm5d* | Y |
| 4. | *Uty* | Y |
| 5. | *Erdr1* | Pseudoautosomal |
| 6. | *Gpr165* | X |
| 7. | *Lrrc10b* | 19 |
| 8. | *Dkk1* | 19 |
| 9. | *Tslp* | 18 |
| 10. | *Pi16* | 17 |
| 11. | *Chodl* | 16 |
| 12. | *Col2a1* | 15 |
| 13. | *Krt75* | 15 |
| 14. | *Fgf9* | 14 |
| 15. | *Gjb6* | 14 |
| 16. | *Thbs4* | 13 |
| 17. | *Hapln1* | 13 |
| 18. | *Adcy2* | 13 |
| 19. | *Dlk1* | 12 |
| 20. | *Syndig1l* | 12 |
| 21. | *Klhl29* | 12 |
| 22. | *2810032G03Rik* | 12 |
| 23. | *Ramp2* | 11 |
| 24. | *Ace* | 11 |
| 25. | *Krt13* | 11 |
| 26. | *Krt14* | 11 |
| 27. | *S100b* | 10 |
| 28. | *Susd2* | 10 |
| 29. | *Bmp5* | 9 |
| 30. | *Stac* | 9 |
| 31. | *1700048O20Rik* | 9 |
| 32. | *Gsta4* | 9 |
| 33. | *Gramd2* | 9 |
| 34. | *1700102P08Rik* | 9 |
| 35. | *Cdh8* | 8 |
| 36. | *Clgn* | 8 |
| 37. | *Lpl* | 8 |
| 38. | *Acan* | 7 |
| 39. | *Scube2* | 7 |
| 40. | *Hif3a* | 7 |
| 41. | *Ntrk3* | 7 |
| 42. | *Gpr27* | 6 |
| 43. | *Klf14* | 6 |
| 44. | *Tril* | 6 |
| 45. | *Eln* | 5 |
| 46. | *Col8a2* | 4 |
| 47. | *Frem1* | 4 |
| 48. | *Col9a2* | 4 |
| 49. | *Ptx3* | 3 |
| 50. | *Agtr1b* | 3 |
| 51. | *Sfrp2* | 3 |
| 52. | *Snord73a* | 3 |
| 53. | *Fam83c* | 2 |
| 54. | *Mylk2* | 2 |
| 55. | *Pcsk2* | 2 |
| 56. | *Adamtsl2* | 2 |
| 57. | *Wnt6* | 1 |
| 58. | *Pappa2* | 1 |
| 59. | *Col9a1* | 1 |
| 60. | *Gsta3* | 1 |
| 61. | *Angptl1* | 1 |
| 62. | *Darc* | 1 |

**Table S3. List of male-lower genes showing ≥1.5-fold lower expression in WT males compared to WT females (P-value <0.05)**

|  | **Genes** | **Chromosome** |
| --- | --- | --- |
| 1. | *Xist* | X |
| 2. | *Eif2s3x* | X |
| 3. | *5530601H04Rik* | X |
| 4. | *Ppef1* | X |
| 5. | *Gda* | 19 |
| 6. | *Malat1* | 19 |
| 7. | *Neat1* | 19 |
| 8. | *4933411K16Rik* | 19 |
| 9. | *Pcdha10* | 18 |
| 10. | *9430076G02Rik* | 18 |
| 11. | *Mc2r* | 18 |
| 12. | *Gm4013* | 18 |
| 13. | *Pcdha9* | 18 |
| 14. | *Abcg1* | 17 |
| 15. | *Rgs11* | 17 |
| 16. | *Xdh* | 17 |
| 17. | *Pisd-ps2* | 17 |
| 18. | *C2* | 17 |
| 19. | *Rhbdl1* | 17 |
| 20. | *Stap2* | 17 |
| 21. | *H2-DMb1* | 17 |
| 22. | *1700071M16Rik* | 17 |
| 23. | *Liph* | 16 |
| 24. | *4930565N06Rik* | 16 |
| 25. | *Nlrc3* | 16 |
| 26. | *Kcnj15* | 16 |
| 27. | *Kng2* | 16 |
| 28. | *Slc7a4* | 16 |
| 29. | *Kifc2* | 15 |
| 30. | *Parp10* | 15 |
| 31. | *Cma2* | 14 |
| 32. | *Ptger2* | 14 |
| 33. | *Gzmd* | 14 |
| 34. | *Gzmc* | 14 |
| 35. | *Mcpt8* | 14 |
| 36. | *Hist1h3d* | 13 |
| 37. | *Fam228b* | 12 |
| 38. | *4930447C04Rik* | 12 |
| 39. | *F730043M19Rik* | 12 |
| 40. | *Ccl8* | 11 |
| 41. | *Doc2b* | 11 |
| 42. | *Pisd-ps1* | 11 |
| 43. | *Rasl10b* | 11 |
| 44. | *Fbxw10* | 11 |
| 45. | *Sfi1* | 11 |
| 46. | *Bzrap1* | 11 |
| 47. | *Cntnap1* | 11 |
| 48. | *Havcr2* | 11 |
| 49. | *Slc16a11* | 11 |
| 50. | *Arg1* | 10 |
| 51. | *Hal* | 10 |
| 52. | *Gstt2* | 10 |
| 53. | *Gm5134* | 10 |
| 54. | *Acp5* | 9 |
| 55. | *Fxyd2* | 9 |
| 56. | *Cxcr6* | 9 |
| 57. | *Mmp3* | 9 |
| 58. | *Cck* | 9 |
| 59. | *Cmtm8* | 9 |
| 60. | *Pstpip1* | 9 |
| 61. | *Mmp13* | 9 |
| 62. | *Mmp10* | 9 |
| 63. | *Hapln4* | 8 |
| 64. | *Chst5* | 8 |
| 65. | *Mboat4* | 8 |
| 66. | *Insl3* | 8 |
| 67. | *Snord68* | 8 |
| 68. | *Acta1* | 8 |
| 69. | *Lrp2bp* | 8 |
| 70. | *Slc27a1* | 8 |
| 71. | *Slc5a5* | 8 |
| 72. | *Nlrc5* | 8 |
| 73. | *Gpr123* | 7 |
| 74. | *Syt17* | 7 |
| 75. | *2810047C21Rik1* | 7 |
| 76. | *Apoc2* | 7 |
| 77. | *Tnfrsf26* | 7 |
| 78. | *Crym* | 7 |
| 79. | *0610005C13Rik* | 7 |
| 80. | *Calcb* | 7 |
| 81. | *Fam19a1* | 6 |
| 82. | *Clec2e* | 6 |
| 83. | *Vwf* | 6 |
| 84. | *B230378P21Rik* | 6 |
| 85. | *Alox5* | 6 |
| 86. | *Spp1* | 5 |
| 87. | *Plac8* | 5 |
| 88. | *Fbxo24* | 5 |
| 89. | *Ppbp* | 5 |
| 90. | *Tfr2* | 5 |
| 91. | *Bmp8b* | 4 |
| 92. | *Cdkn2a* | 4 |
| 93. | *Rex2* | 4 |
| 94. | *Tas1r1* | 4 |
| 95. | *Angptl7* | 4 |
| 96. | *1300002K09Rik* | 4 |
| 97. | *Tnfrsf18* | 4 |
| 98. | *Rnf207* | 4 |
| 99. | *Car9* | 4 |
| 100. | *LOC100038947* | 3 |
| 101. | *Tgm2* | 2 |
| 102. | *Angpt4* | 2 |
| 103. | *Slc52a3* | 2 |
| 104. | *Gm996* | 2 |
| 105. | *1700020A23Rik* | 2 |
| 106. | *Rhov* | 2 |
| 107. | *Siglec1* | 2 |
| 108. | *Olfr1316* | 2 |
| 109. | *Htr2b* | 1 |
| 110. | *Otos* | 1 |
| 111. | *Nphs2* | 1 |
| 112. | *Kcnh1* | 1 |
| 113. | *Fam5c* | 1 |
| 114. | *Creg2* | 1 |

**Table S4. List of PANTHER GO-Slim biological processes revealed by PANTHER overrepresentation test associated with genes specifically dysregulated genes in male MEFs upon HP1γ knockout. Over- or under-representation groups with P<0.05 are presented.**

|  | PANTHER GO-Slim Biological Process | No. of genes in mouse  ref. list | No. of up-regulated genes in male | No. of expected genes | Over(+)/  under(-)  repre-sentation | Fold enrich-ment | P-value |
| --- | --- | --- | --- | --- | --- | --- | --- |
| 1. | regulation of cell cycle (GO:0051726) | 27 | 15 | 4.2 | + | 3.58 | 7.40E-03 |
| 2. | response to interferon-gamma (GO:0034341) | 48 | 21 | 7.46 | + | 2.82 | 7.70E-03 |
| 3. | amino acid transport (GO:0006865) | 63 | 27 | 9.79 | + | 2.76 | 9.35E-04 |
| 4. | DNA replication (GO:0006260) | 152 | 57 | 23.62 | + | 2.41 | 8.09E-07 |
| 5. | antigen processing and presentation (GO:0019882) | 77 | 28 | 11.96 | + | 2.34 | 1.13E-02 |
| 6. | chromosome segregation (GO:0007059) | 84 | 28 | 13.05 | + | 2.15 | 4.66E-02 |
| 7. | DNA repair (GO:0006281) | 168 | 51 | 26.1 | + | 1.95 | 2.11E-03 |
| 8. | B cell mediated immunity (GO:0019724) | 155 | 44 | 24.08 | + | 1.83 | 3.57E-02 |
| 9. | macrophage activation (GO:0042116) | 173 | 49 | 26.88 | + | 1.82 | 1.67E-02 |
| 10. | cell-cell adhesion (GO:0016337) | 362 | 102 | 56.25 | + | 1.81 | 4.43E-06 |
| 11. | DNA metabolic process (GO:0006259) | 352 | 98 | 54.69 | + | 1.79 | 1.44E-05 |
| 12. | receptor-mediated endocytosis (GO:0006898) | 217 | 59 | 33.72 | + | 1.75 | 1.02E-02 |
| 13. | cellular amino acid metabolic process (GO:0006520) | 263 | 69 | 40.86 | + | 1.69 | 7.34E-03 |
| 14. | biological adhesion (GO:0022610) | 577 | 149 | 89.65 | + | 1.66 | 8.32E-07 |
| 15. | cellular defense response (GO:0006968) | 238 | 61 | 36.98 | + | 1.65 | 3.70E-02 |
| 16. | endocytosis (GO:0006897) | 380 | 97 | 59.04 | + | 1.64 | 6.62E-04 |
| 17. | cell adhesion (GO:0007155) | 550 | 140 | 85.46 | + | 1.64 | 5.63E-06 |
| 18. | chromatin organization (GO:0006325) | 244 | 62 | 37.91 | + | 1.64 | 4.12E-02 |
| 19. | immune response (GO:0006955) | 537 | 133 | 83.44 | + | 1.59 | 5.39E-05 |
| 20. | catabolic process (GO:0009056) | 405 | 96 | 62.93 | + | 1.53 | 1.20E-02 |
| 21. | mitosis (GO:0007067) | 365 | 86 | 56.71 | + | 1.52 | 3.42E-02 |
| 22. | cellular component movement (GO:0006928) | 463 | 109 | 71.94 | + | 1.52 | 5.13E-03 |
| 23. | immune system process (GO:0002376) | 1480 | 342 | 229.96 | + | 1.49 | 1.01E-10 |
| 24. | cell cycle (GO:0007049) | 1128 | 248 | 175.26 | + | 1.42 | 1.29E-05 |
| 25. | phosphate-containing compound metabolic process (GO:0006796) | 931 | 203 | 144.66 | + | 1.4 | 3.57E-04 |
| 26. | vesicle-mediated transport (GO:0016192) | 891 | 192 | 138.44 | + | 1.39 | 1.37E-03 |
| 27. | nervous system development (GO:0007399) | 782 | 164 | 121.5 | + | 1.35 | 2.35E-02 |
| 28. | intracellular protein transport (GO:0006886) | 1034 | 216 | 160.66 | + | 1.34 | 2.57E-03 |
| 29. | cellular protein modification process (GO:0006464) | 1341 | 278 | 208.36 | + | 1.33 | 2.54E-04 |
| 30. | system development (GO:0048731) | 1273 | 263 | 197.79 | + | 1.33 | 6.39E-04 |
| 31. | protein transport (GO:0015031) | 1062 | 219 | 165.01 | + | 1.33 | 4.88E-03 |
| 32. | regulation of molecular function (GO:0065009) | 1152 | 231 | 178.99 | + | 1.29 | 1.57E-02 |
| 33. | regulation of catalytic activity (GO:0050790) | 1129 | 226 | 175.42 | + | 1.29 | 2.07E-02 |
| 34. | cell communication (GO:0007154) | 3175 | 635 | 493.32 | + | 1.29 | 3.81E-09 |
| 35. | cellular component organization or biogenesis (GO:0071840) | 1298 | 257 | 201.68 | + | 1.27 | 1.38E-02 |
| 36. | developmental process (GO:0032502) | 2468 | 484 | 383.47 | + | 1.26 | 1.80E-05 |
| 37. | metabolic process (GO:0008152) | 8467 | 1604 | 1315.57 | + | 1.22 | 2.46E-21 |
| 38. | cellular process (GO:0009987) | 7033 | 1330 | 1092.76 | + | 1.22 | 1.95E-15 |
| 39. | primary metabolic process (GO:0044238) | 6997 | 1320 | 1087.17 | + | 1.21 | 6.79E-15 |
| 40. | nucleobase-containing compound metabolic process (GO:0006139) | 3425 | 645 | 532.16 | + | 1.21 | 2.75E-05 |
| 41. | localization (GO:0051179) | 2788 | 523 | 433.19 | + | 1.21 | 8.68E-04 |
| 42. | transport (GO:0006810) | 2658 | 489 | 412.99 | + | 1.18 | 1.21E-02 |
| 43. | response to stimulus (GO:0050896) | 2559 | 469 | 397.61 | + | 1.18 | 2.44E-02 |
| 44. | Unclassified (UNCLASSIFIED) | 9561 | 1120 | 1485.55 | - | 0.75 | 0.00E+00 |
| 45. | response to pheromone (GO:0019236) | 128 | 3 | 19.89 | - | < 0.2 | 7.47E-04 |
| 46. | sensory perception of chemical stimulus (GO:0007606) | 295 | 5 | 45.84 | - | < 0.2 | 4.06E-12 |
| 47. | sensory perception of smell (GO:0007608) | 96 | 1 | 14.92 | - | < 0.2 | 1.14E-03 |

**Table S5. List of PANTHER GO-Slim biological processes revealed by PANTHER overrepresentation test associated with genes specifically dysregulated genes in female MEFs upon HP1γ knockout. Over- or under-representation groups with P<0.05 are presented.**

|  | PANTHER GO-Slim Biological Process | No. of genes in mouse  ref. list | No. of up-regulated genes in male | No. of expected genes | Over(+)/  under(-)  repre-sentation | Fold enrich-ment | P-value |
| --- | --- | --- | --- | --- | --- | --- | --- |
| 1. | cellular glucose homeostasis (GO:0001678) | 62 | 10 | 1.68 | + | 5.95 | 2.28E-03 |
| 2. | skeletal system development (GO:0001501) | 241 | 20 | 6.53 | + | 3.06 | 3.29E-03 |
| 3. | homeostatic process (GO:0042592) | 208 | 17 | 5.64 | + | 3.02 | 1.69E-02 |
| 4. | cell-cell adhesion (GO:0016337) | 362 | 29 | 9.81 | + | 2.96 | 8.61E-05 |
| 5. | ectoderm development (GO:0007398) | 621 | 48 | 16.83 | + | 2.85 | 4.14E-08 |
| 6. | cell adhesion (GO:0007155) | 550 | 42 | 14.91 | + | 2.82 | 8.30E-07 |
| 7. | synaptic transmission (GO:0007268) | 334 | 25 | 9.05 | + | 2.76 | 1.71E-03 |
| 8. | biological adhesion (GO:0022610) | 577 | 43 | 15.64 | + | 2.75 | 1.09E-06 |
| 9. | nervous system development (GO:0007399) | 782 | 55 | 21.2 | + | 2.59 | 5.70E-08 |
| 10. | system development (GO:0048731) | 1273 | 78 | 34.51 | + | 2.26 | 5.93E-09 |
| 11. | cell-cell signaling (GO:0007267) | 598 | 36 | 16.21 | + | 2.22 | 2.43E-03 |
| 12. | mesoderm development (GO:0007498) | 706 | 40 | 19.14 | + | 2.09 | 3.17E-03 |
| 13. | cation transport (GO:0006812) | 579 | 32 | 15.69 | + | 2.04 | 3.51E-02 |
| 14. | ion transport (GO:0006811) | 723 | 37 | 19.6 | + | 1.89 | 4.98E-02 |
| 15. | developmental process (GO:0032502) | 2468 | 118 | 66.9 | + | 1.76 | 1.91E-07 |
| 16. | single-multicellular organism process (GO:0044707) | 1828 | 79 | 49.55 | + | 1.59 | 6.91E-03 |
| 17. | multicellular organismal process (GO:0032501) | 1832 | 79 | 49.66 | + | 1.59 | 7.43E-03 |
| 18. | cell communication (GO:0007154) | 3175 | 134 | 86.06 | + | 1.56 | 2.31E-05 |
| 19. | cellular process (GO:0009987) | 7033 | 234 | 190.63 | + | 1.23 | 2.47E-02 |
| 20. | Unclassified (UNCLASSIFIED) | 9561 | 228 | 259.16 | - | 0.88 | 0.00E+00 |

**Table S6. List of PANTHER GO-Slim biological processes revealed by PANTHER overrepresentation test associated with sexually dimorphic genes. Over- or under-representation groups with P<0.05 are presented.**

|  | PANTHER GO-Slim Biological Process | No. of genes in mouse  ref. list | No. of up-regulated genes in male | No. of expected genes | Over(+)/  under(-)  repre-sentation | Fold enrich-ment | P-value |
| --- | --- | --- | --- | --- | --- | --- | --- |
| 1. | macrophage activation (GO:0042116) | 173 | 7 | 1.16 | + | 6.02 | 4.16E-02 |
| 2. | biological adhesion (GO:0022610) | 577 | 15 | 3.88 | + | 3.87 | 2.06E-03 |
| 3. | immune system process (GO:0002376) | 1480 | 29 | 9.95 | + | 2.92 | 4.08E-05 |
| 4. | system development (GO:0048731) | 1273 | 21 | 8.56 | + | 2.45 | 2.95E-02 |
| 5. | developmental process (GO:0032502) | 2468 | 34 | 16.59 | + | 2.05 | 8.04E-03 |
| 6. | cell communication (GO:0007154) | 3175 | 40 | 21.34 | + | 1.87 | 1.07E-02 |
| 7. | cellular process (GO:0009987) | 7033 | 69 | 47.26 | + | 1.46 | 3.17E-02 |
| 8. | Unclassified (UNCLASSIFIED) | 9561 | 39 | 64.25 | + | 0.61 | 0.00E+00 |

**Table S7. List of sex chromosome encoded gene showing sexually dimorphic expression (P<0.05).**

| Chromo-some | Gene | Log2  (fold change)  WTM vs WTF | P-value | Log2  (fold change)  KOM vs WTM | P-value | Log2  (fold change)  KOF vs WTF | P-value |
| --- | --- | --- | --- | --- | --- | --- | --- |
| Y | ***Eif2s3y*** | 9.42 | 2.83E-183 | 0.22 | 0.0205 | 0.21 | 0.6230 |
| Y | ***Ddx3y*** | 9.85 | 2.67E-180 | 0.09 | 0.5872 | 0.06 | 0.8890 |
| Y | ***Kdm5d*** | 8.43 | 2.87E-130 | 0.06 | 0.6918 | -0.12 | 0.7847 |
| Y | ***Uty*** | 8.33 | 8.46E-109 | 0.26 | 0.1482 | 0.16 | 0.7409 |
| Pseudo-autosomal | ***Erdr1*** | 1.36 | 0.0041 | 0.26 | 0.5854 | 0.58 | 0.2215 |
| X | ***Xist*** | -9.79 | 1.25E-274 | -0.50 | 0.2479 | -0.20 | 0.4298 |
| X | ***Kdm5c*** | -0.55 | 1.30E-07 | -0.21 | 0.0474 | -0.13 | 0.2082 |
| X | ***Eif2s3x*** | -0.66 | 5.94E-05 | -0.29 | 0.0722 | -0.14 | 0.3784 |
| X | ***Ubl4*** | 0.29 | 0.0008 | -0.60 | 2.50E-11 | 0.08 | 0.3449 |
| X | ***Kdm6a*** | -0.52 | 0.0010 | 0.11 | 0.4947 | -0.10 | 0.5100 |
| X | ***5530601H04Rik*** | -0.66 | 0.0022 | 0.06 | 0.7838 | 0.12 | 0.5748 |
| X | ***Htatsf1*** | 0.26 | 0.0074 | -0.59 | 2.13E-09 | -0.02 | 0.8050 |
| X | ***Msl3*** | 0.18 | 0.0078 | -0.20 | 0.0028 | -0.12 | 0.0729 |
| X | ***Smc1a*** | 0.29 | 0.0142 | -0.73 | 6.03E-10 | -0.12 | 0.3113 |
| X | ***2900056M20Rik*** | -0.49 | 0.0174 | 0.76 | 0.0002 | -0.17 | 0.4127 |
| X | ***Tsr2*** | 0.23 | 0.0246 | -0.31 | 0.0032 | 0.17 | 0.0906 |
| X | ***Ftsj1*** | 0.24 | 0.0260 | -0.46 | 2.89E-05 | 0.05 | 0.6219 |
| X | ***Haus7*** | 0.33 | 0.0267 | -0.95 | 3.35E-10 | -0.16 | -0.2988 |
| X | ***Ppef1*** | -1.04 | 0.0399 | 1.779 | 0.0004 | 0.235 | 0.6402 |
| X | ***Uxt*** | 0.23 | 0.0433 | -0.20 | 0.0936 | 0.31 | 0.0058 |
| X | ***Gpr165*** | 1.09 | 0.0444 | -0.65 | 0.2325 | -0.05 | 0.9333 |

**Table S8. Log2 fold change in expression level of *2900056M20Rik and Ppef1* in HP1γ KO males vs WT females.**

| Chromosome | Gene | Log2 (fold change) KOM vs WTF | P-value |
| --- | --- | --- | --- |
| X | ***2900056M20Rik*** | 0.27 | 0.182 |
| X | ***Ppef1*** | 0.74 | 0.1422 |
